# Whi5 is diluted and protein synthesis does not dramatically increase in pre-*Start* G1

**DOI:** 10.1101/2020.06.01.126599

**Authors:** Kurt M. Schmoller, Michael C. Lanz, Jacob Kim, Mardo Koivomagi, Yimiao Qu, Chao Tang, Igor V. Kukhtevich, Robert Schneider, Fabian Rudolf, David F. Moreno, Martí Aldea, Rafael Lucena, Doug Kellogg, Jan M. Skotheim

## Abstract

In their manuscript, Litsios et al.^1^ report a new model for how cell growth and biosynthetic activity control the G1/S transition in budding yeast. In essence, Litsios et al. claim that *Start* is driven by an increasing concentration of the G1 cyclin Cln3 due to a dramatic acceleration of protein synthesis in pre-*Start* G1 and not by the dilution of the cell cycle inhibitor Whi5. While we previously reported that *Start* was in part driven by cell growth during G1 diluting out the *Start* inhibitor Whi5^2^, Litsios et al. report that Whi5 remains at constant concentration during G1, and changes in Whi5 concentration therefore do not contribute to *Start*.

Since Litsios et al. directly contradict several key points of our own model of how cell growth triggers *Start*, we decided to investigate their claims and data. More specifically, we decided to investigate Litsios et al.’s three major claims:

1. Whi5 concentration remains constant during G1
2. Cln3 concentration strongly increases prior to *Start*
3. Global protein synthesis rates increase by 2-3 fold prior to *Start*

We investigated each of these three claims and found that the evidence presented by Litsios et al. does not support their claims due to inadequate analysis methods and flaws in their experiments.

## Whi5 is diluted in G1 in Litsios et al.’s own fluorescence microscopy measurements

We previously observed that Whi5 was diluted during pre-*Start* G1 using wide field fluorescence microscopy. This conclusion is supported by two additional previous observations: First, the expression of *WHI5* is cell cycle dependent and peaks in S/G2^3–5^. Second, Whi5 is a stable protein^2,6^. Thus, whereas the amount of Whi5 molecules is largely stable, cell volume increases during pre-*Start* G1 due to cell growth, resulting in a gradual decrease of Whi5 concentration. This dilution of Whi5 in G1 was also verified using immunoblots (Schmoller et al. shows this in G1 arrested cells^2^; Lucena et al. shows an elutriation experiment with normally sized cells where Whi5/Cln3 decreases (see Fig. S1 for quantitation from this experiment from ^7^). Since Whi5 inhibits *Start* in a dosage-dependent manner, the decrease in Whi5 concentration results in an increase in the rate at which cells progress through *Start*.

The majority of our results and those of Litsios et al. are based on live-cell fluorescence microscopy, which allows observing dilution of Whi5 in asynchronously cycling single cells. After careful subtraction of background fluorescence and cell autofluorescence, the total cellular fluorescence intensity of fluorescently tagged Whi5 can be used as a proxy for the Whi5 amount. Simultaneous estimation of cell volume based on cell segmentations obtained from phase contrast images can then be used to calculate the relative change of Whi5 concentration over time. Since our initial report of Whi5 dilution, several groups have independently reproduced many aspects of our findings in published^8–10^ and in new data reported here that was previously unpublished (Fig. S2–4). Importantly, these verifications of Whi5 dilution were made using different strains, microscopy setups, and image analysis pipelines. In contrast, Litsios et al. concludes that they saw no or very little dilution of Whi5: “While it was recently proposed that the timing of Start is determined by the dilution of Whi5^2^, accumulating evidence from more recent studies contradicts this model. In accordance with our findings, Dorsey et al.^11^ did not observe any dilution of Whi5 under different genetic backgrounds and nutrient conditions, attributing reported changes in the Whi5 concentration to photobleaching.” Importantly, we controlled for photobleaching effects in our original study (cf., Fig. S10 of Schmoller et al., where we show the amount of exposure not to affect the measured Whi5 dynamics^2^). Moreover, using the same imaging conditions and fluorophore mCitrine, we found that all other proteins we examined (Swi4, Bck2, Whi3) did not show similar dilution^2^. We cannot comment further here on the work of Dorsey et al. because they used a complex confocal-based sN&B imaging approach that we do not have expertise with.

Due to the discrepancy between our observation and Litsios et al.’s conclusion that Whi5 shows little or no dilution, we sought to examine their data more carefully, which indicated that the concentration of Whi5 was constant during G1 (cf., Litsios et al Fig. 5a^1^). Upon examination of the individual Whi5-mCherry concentration traces published in Litsios et al., we were surprised to see that Whi5 concentration decreased in 44 of 49 cells from birth to *Start* (Fig. 1a). To draw the conclusion that Whi5 concentration was constant from their microscopy data, Litsios et al. performed a rather unusual normalization and averaging procedure. They normalized the Whi5 concentration of each individual cell to the time of cell birth, and then aligned all single-cell traces at *Start* to calculate the time-evolution of the average shown in their Fig. 5a - which indeed shows only a modest decrease in Whi5 concentration. However, given the large variability of pre-*Start*-G1 durations, aligning cell traces at *Start* while simultaneously normalizing to the concentration at birth largely masks the underlying dynamics and is completely inadequate to determine whether or not Whi5 is diluted in single cells. To make this point absolutely clear, consider 4 cells that all grow with similar exponential growth rate, but show varying durations of the period from birth to *Start* (Fig. 1b). Assuming ideal dilution of a stable pool of Whi5, the Whi5 concentration in each cell will then decrease in inverse proportion to cell volume as cells are growing during G1. If normalized to the initial concentration at birth, each cell will then show a similar decrease over time. Thus, even though each single-cell trace will have different length, the mean of the traces aligned at birth will accurately represent the average dynamics of Whi5 dilution (Fig. 1c). If we instead align all normalized traces at *Start*, as Litsios et al. did, it becomes immediately obvious that the resulting mean does not reflect the dynamics of dilution, but instead strongly depends on the distribution of pre-*Start*-G1 duration (Fig. 1d).

**Figure 1:**
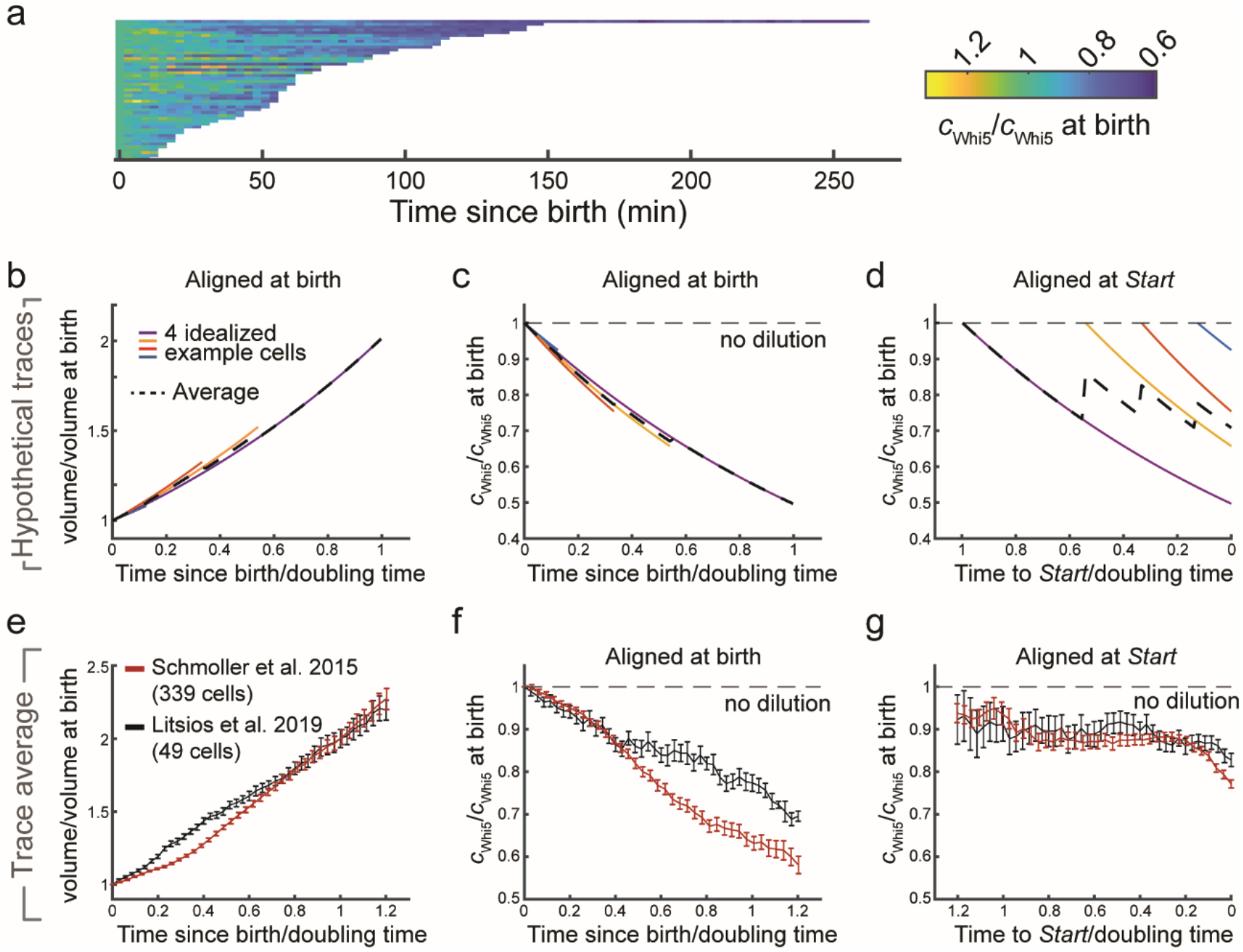
Whi5 is diluted during pre-Start G1. **a)** Single cell traces of Whi5-mCherry concentration reported by Litsios et al. are shown over time from cell birth to *Start*. Each trace is normalized on the initial value at cell birth. n=49 cells. **b-d)** Illustration of the effect of single-cell normalization and alignment on the observed apparent dynamics of Whi5 dilution. We assume 4 idealized cells with different pre-*Start* G1 duration, each growing exponentially with similar growth rate (b, volume doubling time is estimated from the mean behavior) and diluting a stable pool of Whi5. The mean accurately reflects the single-cell dilution if concentration traces are normalized and aligned at cell birth (c). If single-cell-traces are normalized at birth and aligned at *Start* instead, the mean does not reflect the single-cell dynamics and strongly depends on the distribution of pre-*Start*-G1 duration (d). **e-g)** Data by Litsios et al. on Whi5-mCherrry^1^ and Schmoller et al. on Whi5-mCitrine^2^ show similar Whi5 dilution if plotted accurately. Since the data sets were obtained using different growth media, we first determined the volume doubling time from the mean relative volume growth over time (e). If normalized and aligned at birth, both data sets show dilution of Whi5 as cells progress through pre-*Start* G1 (f). If normalized at birth and aligned at *Start*, dilution is largely masked in the mean behavior (g).

Given these problems associated with normalizing single-cell traces at one time point, *i.e*. cell birth, and aligning them at another, *i.e. Start*, we wondered whether the fact that Litsios et al. did not observe Whi5 dilution may be simply an artifact due to their unusual alignment and normalization procedure. We therefore replotted the raw data of Whi5-mCherry concentration shown in Litsios et al., and found that Whi5 is clearly diluted during G1, with dynamics that are comparable to those we previously observed^2^ (Fig. 1 e,f). We note that we have rescaled time with the amount of time it takes a cell to double in volume in the two different media conditions (based on the averages shown in Fig. 1e). In contrast, if we align our data at *Start* while normalizing at birth, Whi5-dilution is masked due to inadequate averaging (Fig. 1g).

In summary, after a correct analysis, the microscopy data shown in Litsios et al. show clear dilution of Whi5 and are qualitatively consistent with the data of Schmoller et al. and several other groups. Whether the smaller quantitative difference in the dilution behavior observed in the two data sets reflects a strain- or condition-dependent difference or is due to different image analysis protocols is unclear. To clarify this, we requested their raw image files, but did not receive them.

## Mass spectrometry data from Litsios *et al*. are inconclusive

As an approach orthogonal to fluorescent microscopy, the authors employed mass spectrometry to determine relative changes in Cln3 and Whi5 protein concentrations as cells that were initially in G1 progressed through the cell cycle. They used centrifugal elutriation to enrich for pre-*Start* cells and monitored changes in the relative abundance of Whi5 and Cln3 peptides as these semi-synchronous cultures progressed through the cell cycle. Likely because the abundance of Whi5 and Cln3 peptides are too low to detect using conventional Data-dependent acquisition (DDA) proteomics, the authors used parallel reaction monitoring (PRM) to quantify the relative abundance of two Whi5 and two Cln3 peptides.

The authors performed 4 independent elutriation time courses and claim that their quantification of Whi5 and Cln3 peptides via targeted proteomics supports their measurements of Whi5 and Cln3 made using microscopy. However, we reviewed their proteomic analysis (see methods, Fig. S5, and Table S1 for a complete description of our re-analysis) and found that it is complicated by a striking inconsistency in the content of their protein samples. Despite a relatively consistent injection of bulk peptide material into their LC-MS (Fig. 2a, as inferred by the approximate total ion current (TIC) from Litsios et al.’s raw data), the peptide peak areas of several common yeast housekeeping proteins greatly vary from run-to-run (one example is shown in Fig. 2b; similar plots for all experiments are shown in supporting Fig. S5). This suggests that the proportion of peptides originating from yeast proteins varies greatly from sample-to-sample and that, in some cases, nearly all the injected mixture is composed of foreign peptides due to contamination. Though not acknowledged in their manuscript, the authors were aware of this contamination (personal correspondence). They suspected that the yeast pellets obtained from the elutriation time course were insufficiently washed and that the source of the contaminating peptides was remnant peptone from the YPD yeast media. Because proteins in peptone are animal in origin, the authors attempted to correct for the suppressive presence of animal peptides by applying a normalization to a set of yeast housekeeping proteins (like those depicted in Fig. 2b).

**Figure 2:**
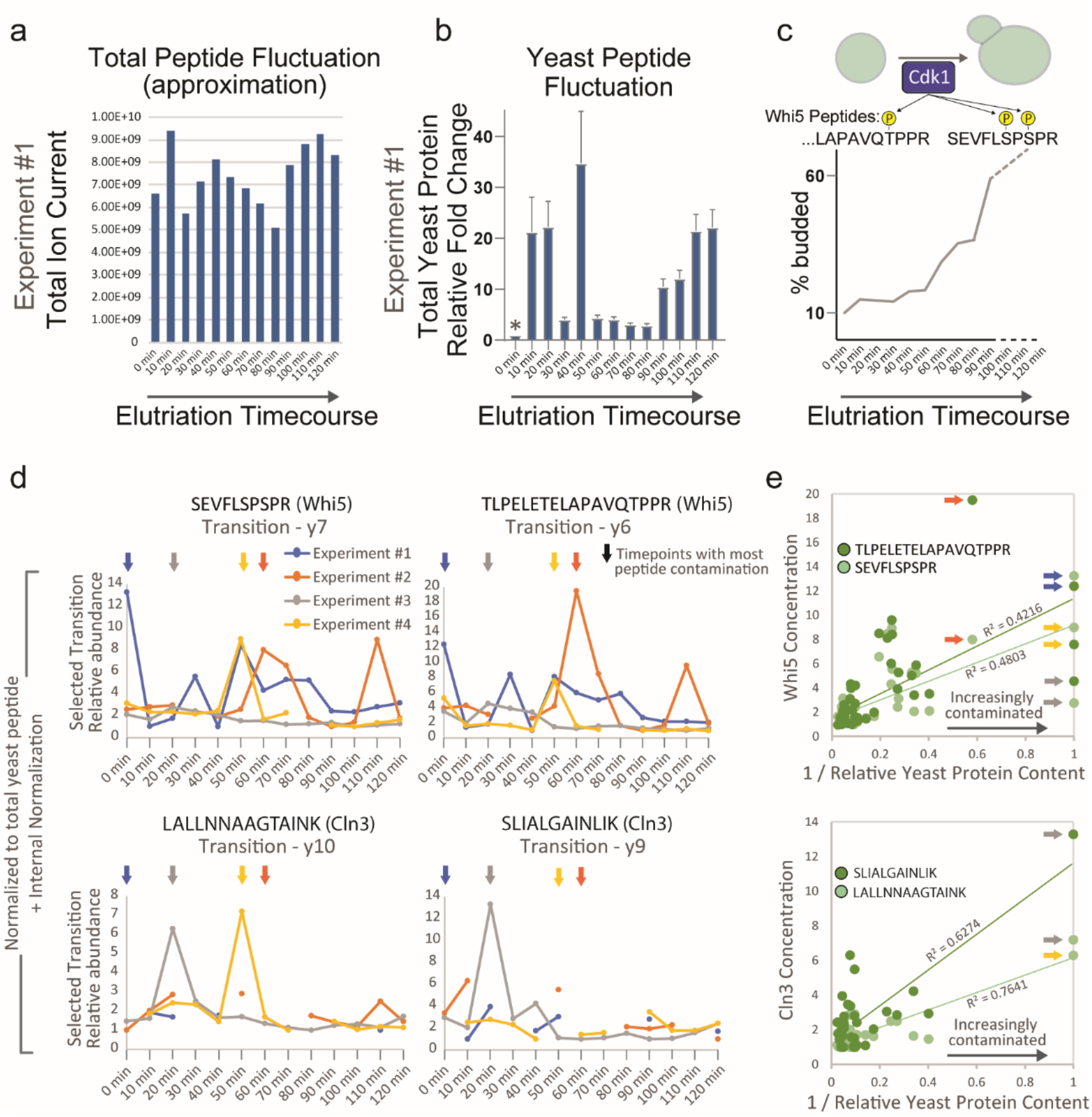
Mass spectrometry data is inconclusive. **a)** Total peptide load estimated for each time point in the elutriation time course based on the measurement of total ion current (TIC). Experiment #1 is shown as an example (see supplemental Fig. S5 for all replicate experiments). **b)** Total yeast peptide amount based on the measurement of 6 yeast housekeeping peptides. Date from experiment #1 is shown as an example. MS1 peak areas of peptides from Enolase, GAPDH, and ADH1 were used to estimate changes to the total amount of yeast protein. Yeast peptide fold change was calculated relative to the time point with the least amount of yeast peptide signal (denoted by asterisk). These fold changes were used to normalize the PRM measurements of Whi5 and Cln3 peptides. Error bars represent SEM. **c)** Top: Schematic of Whi5 peptides indicating known phosphorylation sites of Cdk1. Bottom: Bud index from Litsios et al. indicating the degree of cell cycle synchrony and progression through the cell cycle. Whi5 is highly phosphorylated at the G1/S transition (see Fig. S6). **d)** PRM measurements (normalized to the amount of total yeast protein) for Cln3 and Whi5 peptides throughout the elutriation time course. All four experiments are plotted. Litsios *etal*. used the single, most intense transition to quantify the elution curves for Whi5 and Cln3 peptides. We used the same ions in our re-analysis. Missing data points represent PRM measurements that did not pass a quality metric designated by the authors (Qvalue > 0.01). Arrows denote the time points in each of the 4 replicate time courses that contained the largest amount of foreign peptide contamination (*i.e*., the least amount of yeast peptide). Table S1 contains a step-by-step derivation of the plotted datapoints. **e)** The concentration calculated for each Whi5 and Cln3 peptide throughout all experimental time courses (*i.e*., the data points from (d)) plotted against the extent of peptide contamination in each of those timepoints. The extent of foreign peptide contamination is inferred from the relative amount of yeast peptides in each sample (plotted in (b) and S5), which was also used in the determination of relative concentration. Arrows correspond to the timepoints in (d). The most contaminated sample in Experiment #2 was manually excluded by Litsios et al., thus the orange arrow points to the second most contaminated timepoint. The arrows that are absent in the Cln3 plot correspond to timepoints where the PRM measurement of Cln3 did not pass the required quality metric and were excluded.

We think the following points demonstrate this proteomic dataset is not suited to measure the concentration changes of Cln3 and Whi5 through the cell cycle. **1)** The extreme extent of the MS1-level normalization means that the Whi5 and Cln3 PRM signals are regularly adjusted by over 20fold, even between adjacent time points (Fig. 2b, S5). **2)** Both Whi5 peptides monitored by the PRM analysis contain phosphosites targeted by the cyclin-dependent kinase Cdk1 at *Start*, which lead to Whi5’s nuclear export^12–15^. The stoichiometry of Whi5 phosphorylation after Start is high and can be inferred from its phospho-dependent shift in a phostag gel (Fig. S6). Thus, the concentration of non-phosphorylated forms of these peptides, which is what the PRM analysis measures, should decrease significantly when cells proceed through *Start* and are in S phase (see peptide schematic in Fig. 2c showing Cdk sites and bud index throughout elutriation time course). That Litsios et al. do not observe a decrease in the concentration of unphosphorylated Whi5 peptides through the cell cycle suggests that their measurements do not reflect *in vivo* changes in Whi5 concentration. **3)** Though the authors reported that four biological replicates were used to calculate the changes in Cln3 concentration, many of the data points surrounding *Start* (the period in which they see a “pulse”) were excluded based on a PRM quality control metric (Fig. 2d, Table S1). **4)** When the experiments are plotted separately, the noisy nature of the data is more evident, and it is clear that the Cln3 “pulse” reported at t=20min is primarily driven by experiment #3 and is less apparent in the other experiments. We also note that the Whi5 time courses contain many non-reproducible “pulses” in concentration. The authors of Litsios et al. confirmed the correspondence of our re-analysis with the data they used to generate the plots in Figure 4e of their manuscript. **5)** When considering all their experiments together, nearly every peak in concentration corresponds to the timepoints within each time course that contain the greatest extent of peptide contamination (see arrows in Fig. 2d), which are therefore subjected to the most adjustment during normalization. In fact, when all the timepoints from all time courses are plotted together, the derived Whi5 concentrations correlate with the extent of peptide contamination (Fig. 2e). Cln3 exhibits a similar, though less visually dramatic correlation (Fig. 2e), which could be explained by the fact that many of the most contaminated Cln3 timepoints are excluded by the Qvalue (some of these timepoints have no detectable Cln3 PRM signal). This observation indicates that the ‘normalization’ used to derive Cln3 and Whi5 concentration by Litsios et al. is significantly biased by the extent of contamination in each of their peptide samples.

Given these caveats, we do not think that the presented mass-spectrometry data support the claim that Cln3 concentration sharply increases in G1 prior to Start nor the claim that Whi5 is not diluted. Importantly, since the microscopy data presented by Litsios et al. show a clear dilution of Whi5 when aligned correctly (Fig. 1), the mass-spectrometry data is not consistent with the microscopy data presented in the same study. Moreover, similar elutriation experiments were performed independently by two other research groups, both of which found that Cln3 concentrations were largely constant or slightly increasing, and that Whi5 was diluted during G1^7,10^.

## There is no dramatic increase in protein synthesis rates leading up to *Start*

A central part of the model suggested by Litsios et al. is that protein synthesis rates increase 2-3 fold in G1 leading up to *Start*. This is important because this increase in the protein synthesis rate would drive concentration changes for highly unstable proteins, such as the cell cycle activator Cln3. Should the authors’ interpretation be correct, then there would be a 2-3 fold increase in the Cln3 concentration leading up to *Start* that triggers this cell cycle transition. Given the problems of the mass spectrometry data described above, we were curious whether the microscopy data support this claim better.

To begin our reanalysis of Litsios et al.’s claim, we outline why unstable proteins should be sensitive to changes in protein synthesis rates. The change in the amount of an unstable protein, p, synthesized at a constant rate, s, and degraded in a first-order reaction at a rate d, is given by dp/dt = s - d p. Since we are interested in rapidly degraded proteins, the concentrations will reach steady state so that p = s/d, *i.e*., the amount of the unstable protein is directly proportional to the protein synthesis rate. Then, the concentration of the unstable protein in a cell of volume v is [p] = s / (d v) (we note that the nuclear volume, where these proteins reside, is approximately proportional to the cell volume^16,17^).

To support the claim that Cln3 concentration increases several-fold leading up to *Start*, Litsios et al. present fluorescence microscopy data of cells expressing *CLN3-2A-sfGFP*. sfGFP is separated from Cln3 upon translation through the viral self-cleaving 2A sequence, which enables imaging despite the fact that Cln3 is very unstable. The total amount of sfGFP in each cell at each time point is then calculated by multiplying the mean pixel intensity by the estimated area. Then, the Cln3 synthesis rate is calculated by taking the first derivative of a Gaussian Process smoothing fit of the total GFP curve. Since Cln3 synthesis rate is proportional to Cln3 abundance, they estimate a ~2-fold increase in Cln3 abundance (not concentration) around *Start*.

While protein amounts may be important, protein concentrations are typically the more relevant quantities for kinase reactions such as those that may be driven by Cln3-Cdk1. When we convert the Litsios et al. Cln3 abundance measurements to concentration measurements, the result is a less dramatic increase (Fig. 3a). Rather than a multiple fold-increase, we see a ~50% increase over the 40 minutes preceding *Start*, which is certainly a sizable effect. However, when we plotted the Litsios et al. data over a slightly larger range than what was shown in their publication (the data shown in their Fig. 5c are truncated), we see that already 80 minutes before *Start*, Cln3 is estimated to be at a high concentration that was comparable to the concentration at *Start*, casting some doubt on the conclusion that Cln3 concentration dramatically increases from cell birth to *Start*. Given that Gaussian Process smoothing effectively averages the signal over multiple time points, we thought that some decrease of the protein synthesis rates around budding may have been due to the authors not including fluorescence from the bud, but only the mother cell body. To shed light on this issue, we requested the raw microscopy files associated with the measurements of Litsios et al. multiple times from the authors, but we did not receive any. Eventually, Dr. Heinemann was willing to share some of the requested raw imaging data, but only if we would sign a highly restrictive, unusual data transfer agreement (included in the supporting material). We declined to sign the agreement and were therefore not given access to any of the unprocessed imaging data.

**Figure 3:**
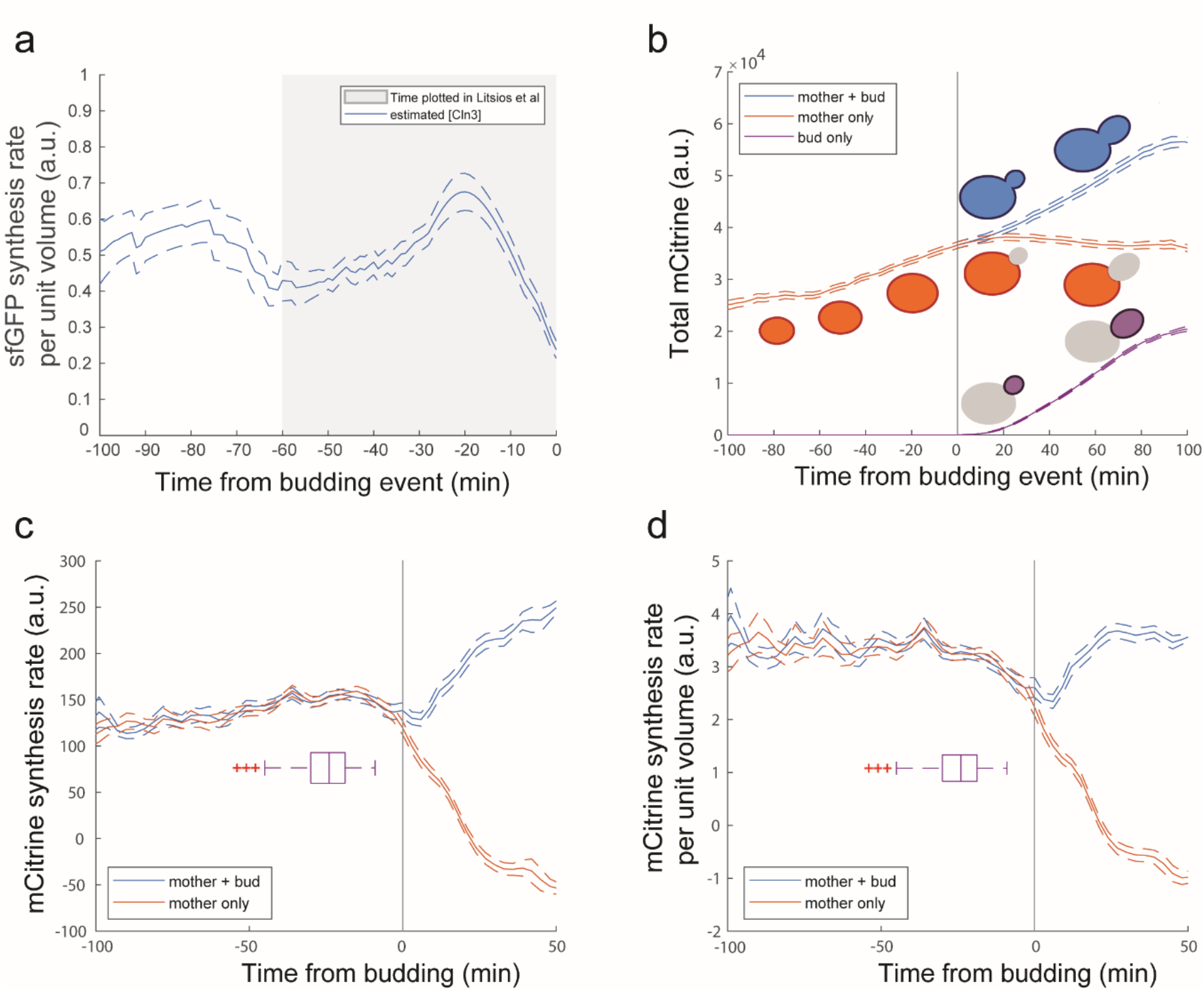
Protein synthesis rates do not dramatically increase prior to *Start*. **a)** Estimate of the Cln3 concentration from Litsios et al.’s data. Traces were aligned to budding and the sfGFP synthesis rate and volume data for 41 *CLN3-2A-sfGFP* cells was first used to estimate the sfGFP synthesis rate as shown in Litsios et al Fig. 5c, then this estimate was divided by the volume of each cell to estimate the Cln3 concentration. The volume dynamics for each cell was estimated using a linear fit. This results in an estimate of the Cln3 concentration at each time point (see text). The solid line represents the mean value and the dashed lines represent the standard error at each time point. The gray background represents the time axis plotted in Litsios et al. Fig. 5c, while here we show a more complete time range. **b)** Mean cellular mCitrine dynamics expressed from an integrated *ACT1pr-mCitrine* allele. Data are shown for the whole yeast, mother cell body only, and for the bud only. The fluorescence microscopy data of 163 *ACT1pr-mCitrine* cells was from Chandler-Brown et al.^18^. Each trace was aligned to the time of budding (black vertical line). The solid lines represent the mean value and the dashed lines represent the standard error at each time point. **c)** Mean cellular mCitrine synthesis rate. Data from b) was smoothed using a Gaussian process regression function using a similar approach to that used by Litsios et al. Then, the rate was calculated by subtracting the adjacent time point and dividing by the 3 minutes time between movie frames. Each trace was aligned to the time of the budding (black vertical line). The solid lines represent the mean value and the dashed lines represent the standard error at each time point. The boxplot shows the distribution of time points at which each cell passes *Start* (Whi5 nuclear exit). **d)** The concentration of unstable proteins such as Cln3 was estimated as the rate of mCitrine synthesis per unit volume in *ACT1pr-mCitrine* cells (see text). Data for individual mCitrine synthesis rates from c) was divided by the volume at each time point. Each trace was aligned to the time of the budding (black vertical line). The solid lines represent the mean value and the dashed lines represent the standard error at each time point. The boxplot shows the distribution of time points at which each cell passes *Start* (Whi5 nuclear exit).

While we did not have access to the raw images to analyze data from *CLN3-2A-sfGFP* expressing cells, the increase in Cln3 was proposed to reflect a global increase in protein synthesis rates and should therefore be visible in the production rates of most proteins. Supporting this view, Litsios et al. also examined sfGFP expressed from a constitutive *TEF1* promoter and found a similar increase in protein synthesis as they found for their Cln3 reporter. This led us to reanalyze some of our own data on cells expressing mCitrine fluorescent protein from an *ACT1* promoter^18^. We observe that the total cellular fluorescence in the mother body increases smoothly up until bud emergence, following which cell growth is due to increases in the bud volume (Fig. 3b; Fig. S7). We can then use these fluorescence traces to estimate the protein synthesis rate (Fig. 3c) and then divide by the cell volume to estimate the concentration of unstable proteins including Cln3 (Fig. 3d). Unlike Litsios et al., we estimate a much more constant concentration of unstable proteins, like Cln3, prior to *Start*. Following *Start* and around bud emergence, we see a slight decrease in the protein synthesis rate, which is followed by an increase to a similar protein synthesis rate per unit mass as before *Start*. We see no evidence of a 2-3 fold increase in the global protein synthesis rate. These results are consistent with our previous results examining a stabilized Cln3 protein expressed from the endogenous *CLN3* locus^2^. Moreover, we note that our results are also consistent with bulk radiolabeling experiments looking at total protein synthesis rates in cells synchronized by elutriation that did not find any changes in global synthesis rate^19^. Thus, while there may be some specific proteins whose abundance increases through G1, this is unlikely to reflect changes in the global protein synthesis rate prior to *Start*.

## Concluding remarks

The revised model of *Start* proposed by Litsios et al. rests on two important notions. First, that Whi5 is not diluted. Second, that there is a 2-3 fold increase in protein synthesis rates that drive a pulse of Cln3 synthesis to drive *Start*. Having reanalyzed the data from Litsios et al, we see no or at best very limited evidence of either of these claims due to both experimental limitations and poor analysis methods. Although one other study recently reported a constant concentration of Whi5 through G1^11^, this conclusion is now quite isolated. At this point, we are aware of at least 9 independent research groups (including the authors) that have corroborated Whi5 dilution. These studies used both immunoblots and fluorescence microscopy, and were done using different strain backgrounds, microscopes, and analysis methods. Importantly, our reanalysis here of Litsios et al.’s data shows that Whi5 is diluted during pre-*Start* G1 in this data set as well. On average, Whi5 concentration decreases by 20% prior to *Start*, which is consistent with the average amount of cell growth during G1 in these conditions. As discussed in a recent paper by Qu et al., the relative amount of Whi5 dilution increases as cell growth rates decrease, and G1 durations increase, in poorer nutrient conditions^8^. Finally, it is important to note that our claim is that Whi5 dilution is one mechanism through which cell growth in G1 promotes cell cycle progression. We do not claim Whi5 dilution is the only such mechanism, which is consistent with a recent study proposing that cell growth triggers relative concentration changes of a broader group of cell cycle regulators to drive *Start*^10^.

## Supplementary Figures

**Figure S1:**
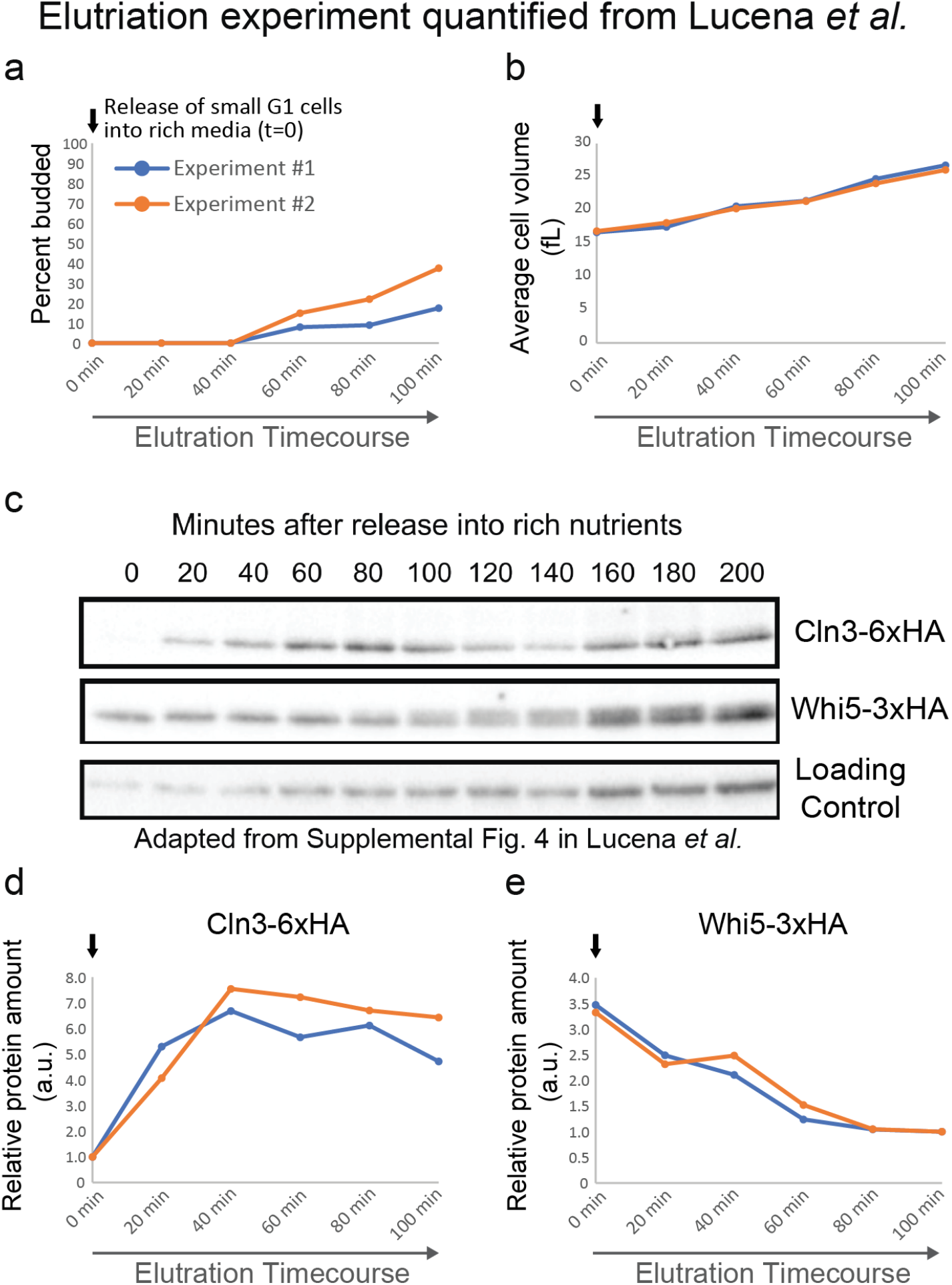
Elutriation experiment performed by the Kellogg lab. Immunoblot was adapted from Lucena *et al*. and quantified here with bud index and cell size measurements. Cells were grown in poor carbon conditions to facilitate the isolation of small G1 cells during elutriation. After isolation via elutriation, the small G1 cells were released into glucose rich media (t=0). Two independent experiments were performed. **a)** Budding index and **b)** Average cell volume were quantified throughout the time course. **c)** Whi5 and Cln3 abundance was quantified via western blotting and normalized to a loading control. The results obtained from 2 independent experiments are plotted over time in **d)** and **e)**. *Start* takes place at the end of the time course. Cln3 concentration increases as cells initiate growth in the rich media, but then flattens out prior to *Start*. Whi5 is diluted in pre-*Start* G1 as described in Schmoller *et al*. and consistent with the additional fluorescence measurements reported here.

**Figure S2:**
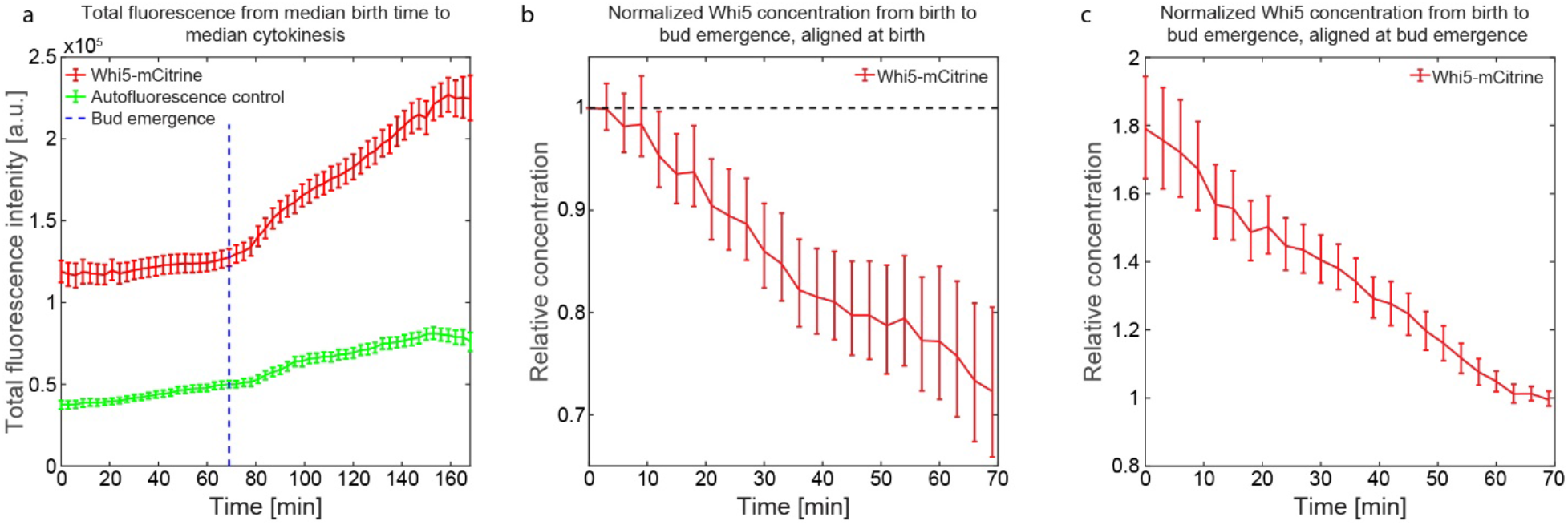
Whi5 concentration dynamics in G1 measured using fluorescence microscopy as measured by the Schneider lab. **a)** Mean total fluorescent intensity over the cell cycle for *WHI5-mCitrine* (N=100) and wild-type cells (N=59) are shown. **b-c)** Relative concentration dynamics for *WHI5-mCitrine* defined as mean fluorescence intensity after subtraction of the wild-type autofluorescence divided by the cell volume and then normalized to the value at birth (b) or to the value at bud emergence (c). Cell traces are aligned at either birth (b) or bud emergence (c). Cells were grown in SCGE as described by Schmoller et al.^2^. Whiskers denote 95% confidence intervals, which were determined from 50000 bootstrap samples.

**Figure S3:**
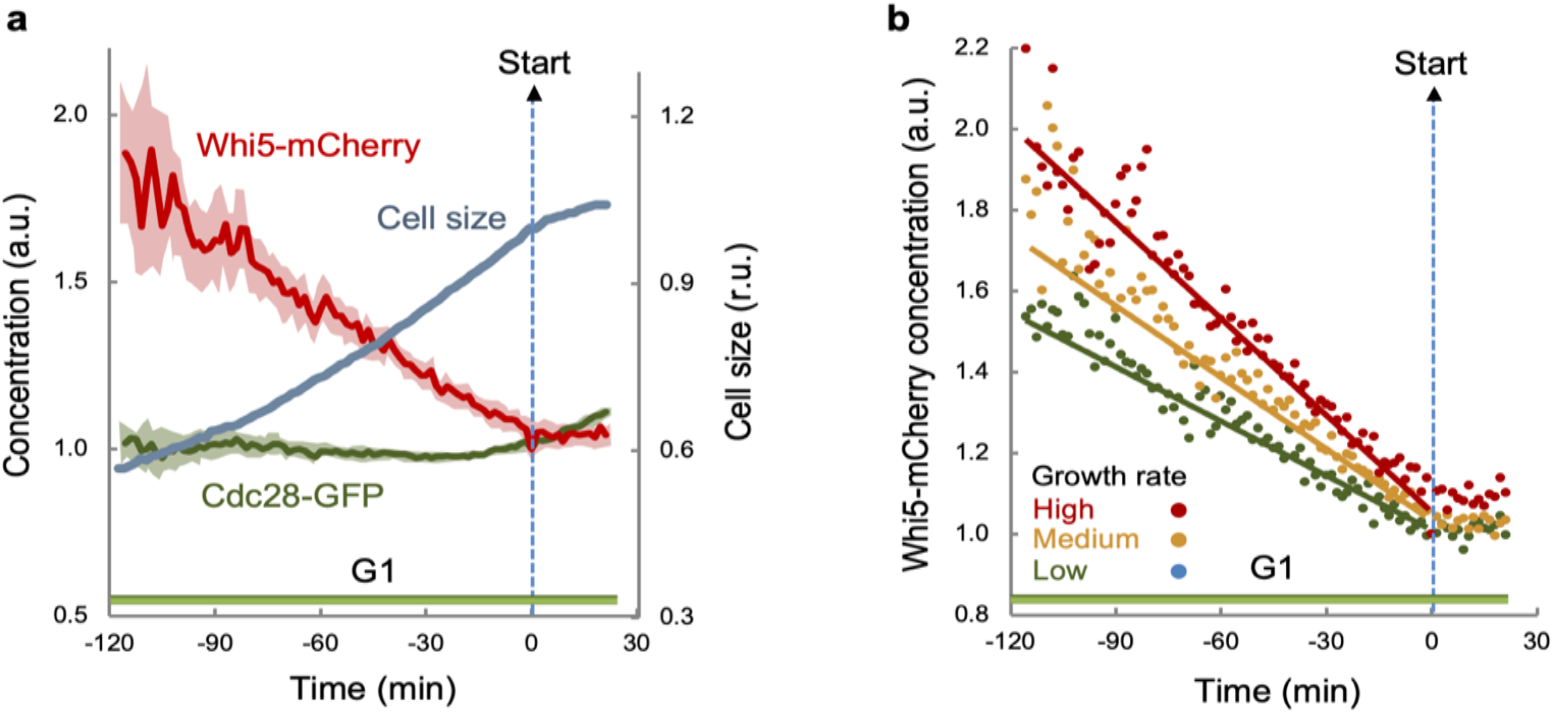
Whi5 dilution in G1 scales with growth rate as measured by the Aldea lab. **a**) Cells expressing Cdc28-GFP and Whi5-mCherry were analyzed by time-lapse microscopy during G1 progression and entry into the cell cycle, and time-lapse data from individual cells were aligned to *Start*. Mean (N>100) relative cellular concentrations of Cdc28-GFP (green) and Whi5-mCherry (red), and the respective confidence limits (α=0.05) for the mean are plotted. Relative mean cell volumes (blue) are also shown. Concentrations were normalized to their value at *Start*. **b**) Cells expressing Whi5-mCherry were analyzed by time-lapse microscopy during G1 progression and entry into the cell cycle. Time-lapse data from individual cells were aligned to *Start* and binned into three groups by growth rate in G1 (high: red; medium: yellow; low: green). Mean (N>50) cellular concentrations of Whi5-mCherry are plotted as a function of time during G1 and normalized to their values at *Start*.

**Figure S4:**
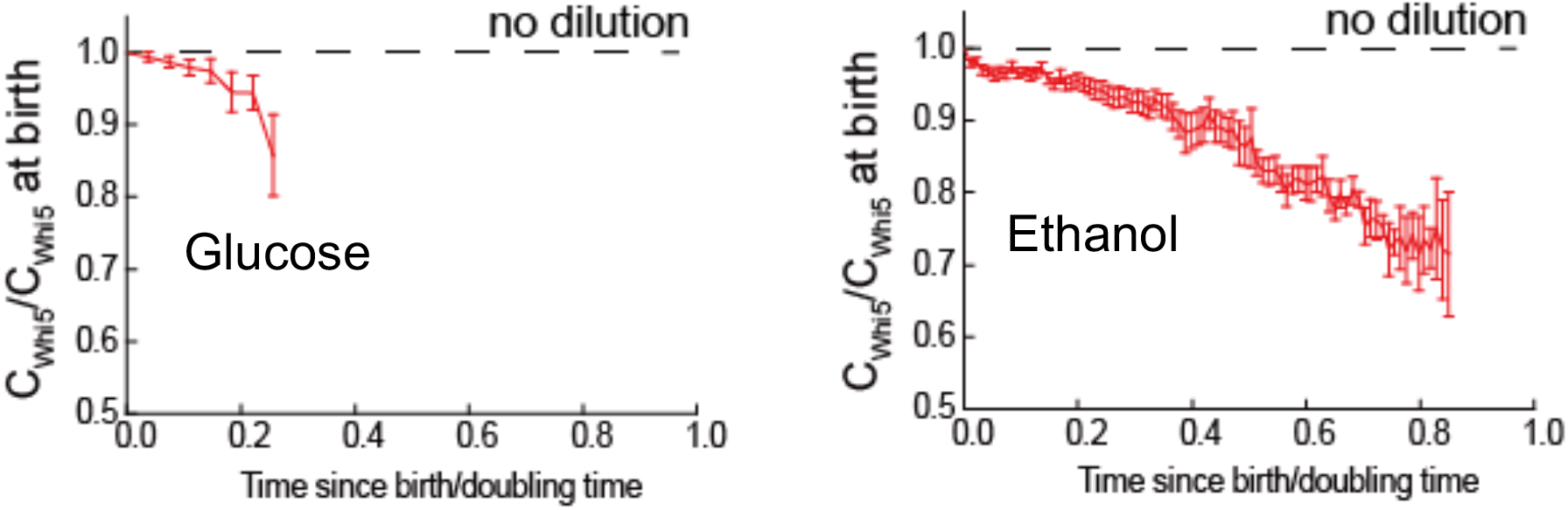
Whi5 dilution in G1 scales with growth rate measured by the Tang lab. Single daughter cell traces of Whi5-tdTomato concentration were aligned and normalized at birth and the cells were followed from birth to *Start*. See Qu. et al.^8^ for details of growth conditions and segmentation methods (the doubling time is 82±1 minutes and n = 73 in glucose media, and the doubling time is 286±9 minutes and n = 83 in ethanol media; we note that for the data presented here, Qu et al. quantified total cellular fluorescence and cell volume following Schmoller et al.^2^). Birth is defined as the time point when Whi5 enters the nucleus and *Start* is defined as the time point when Whi5 exits the nucleus. Data are shown as mean ± SEM.

**Figure S5:**
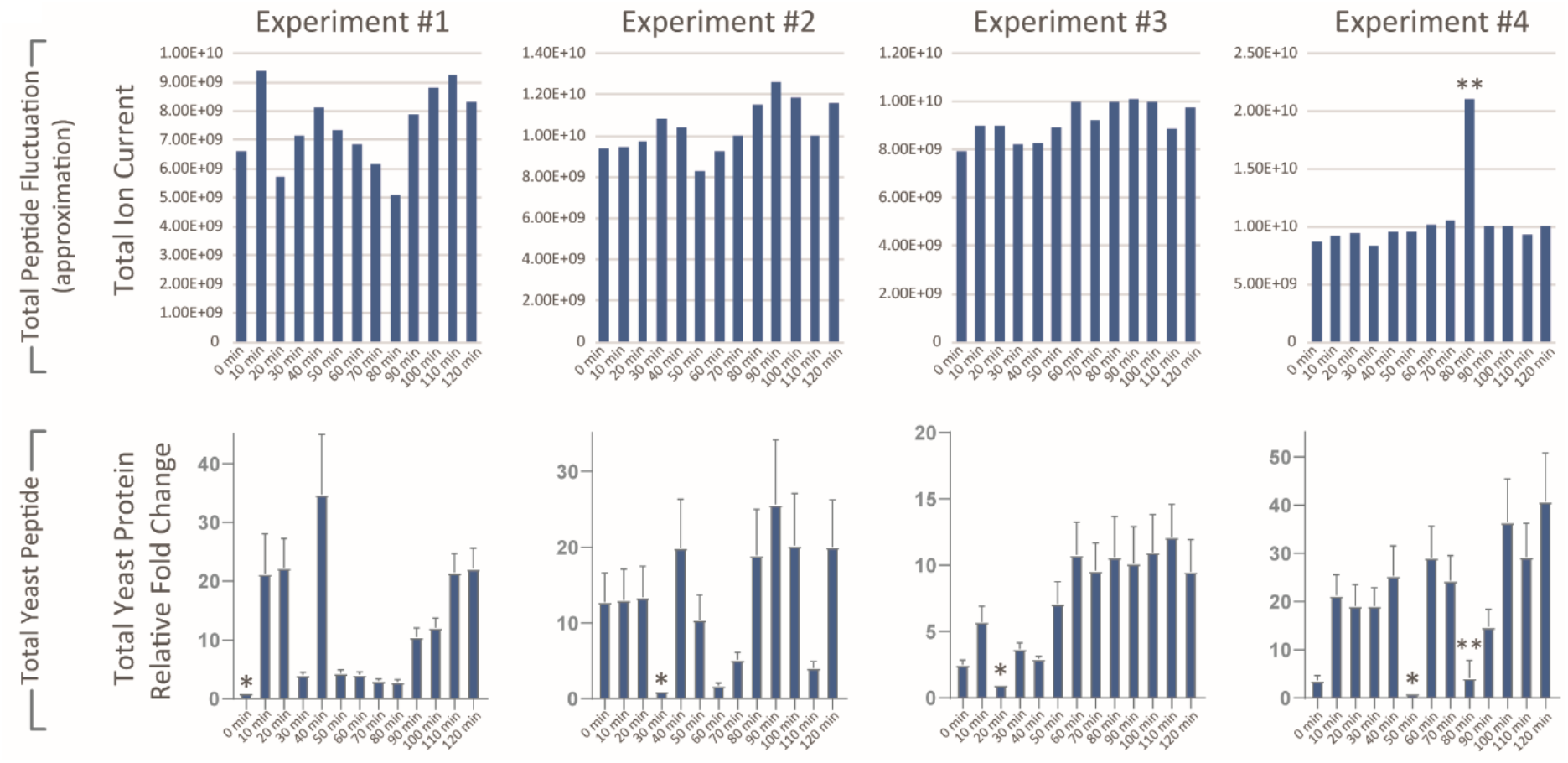
Supporting analysis of the mass spectrometry data shown in Fig. 2. Top row: Total peptide load estimated for each time point in the elutriation time course based on the measurement of total ion current (TIC). Bottom row: Total yeast peptide amount based on the measurement of 6 yeast housekeeping peptides. Date from experiment #1 is shown as an example. MS1 peak areas of peptides from Enolase, GAPDH, and ADH1 were used to estimate changes to the total amount of yeast protein. Yeast peptide fold change was calculated relative to the time point with the least amount of yeast peptide signal (denoted by asterisk). These fold changes were used to normalize the PRM measurements of Whi5 and Cln3 peptides. Double asterisk in experiment 4 denotes a run with compromised peptide chromatography (this data point was removed from all analyses).

**Figure S6:**
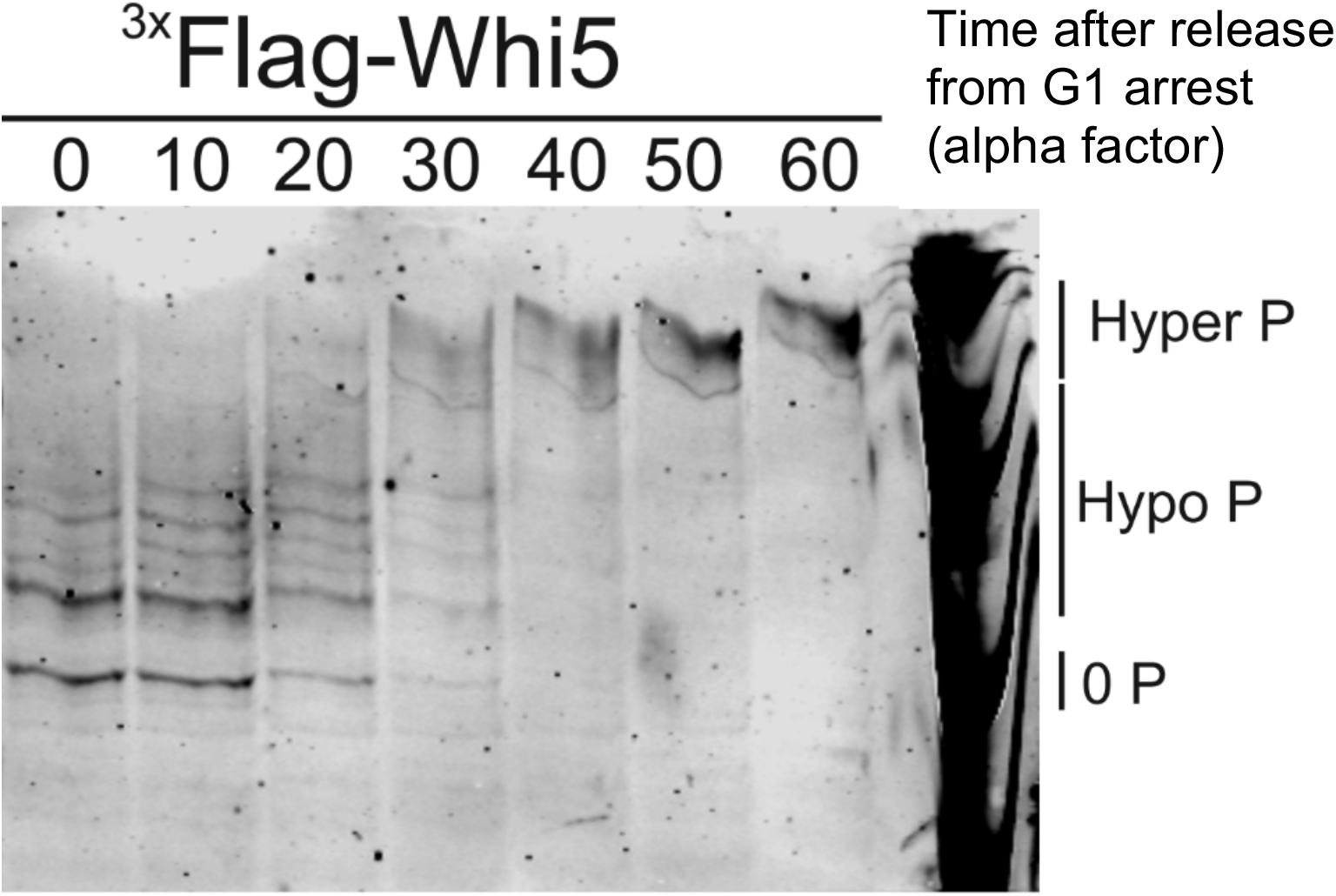
Phos-tag immunoblot time course measuring distinct hypo- and hyperphosphorylated isoforms of 3xFlag-Whi5 after release from alpha factor G1 arrest.

**Figure S7:**
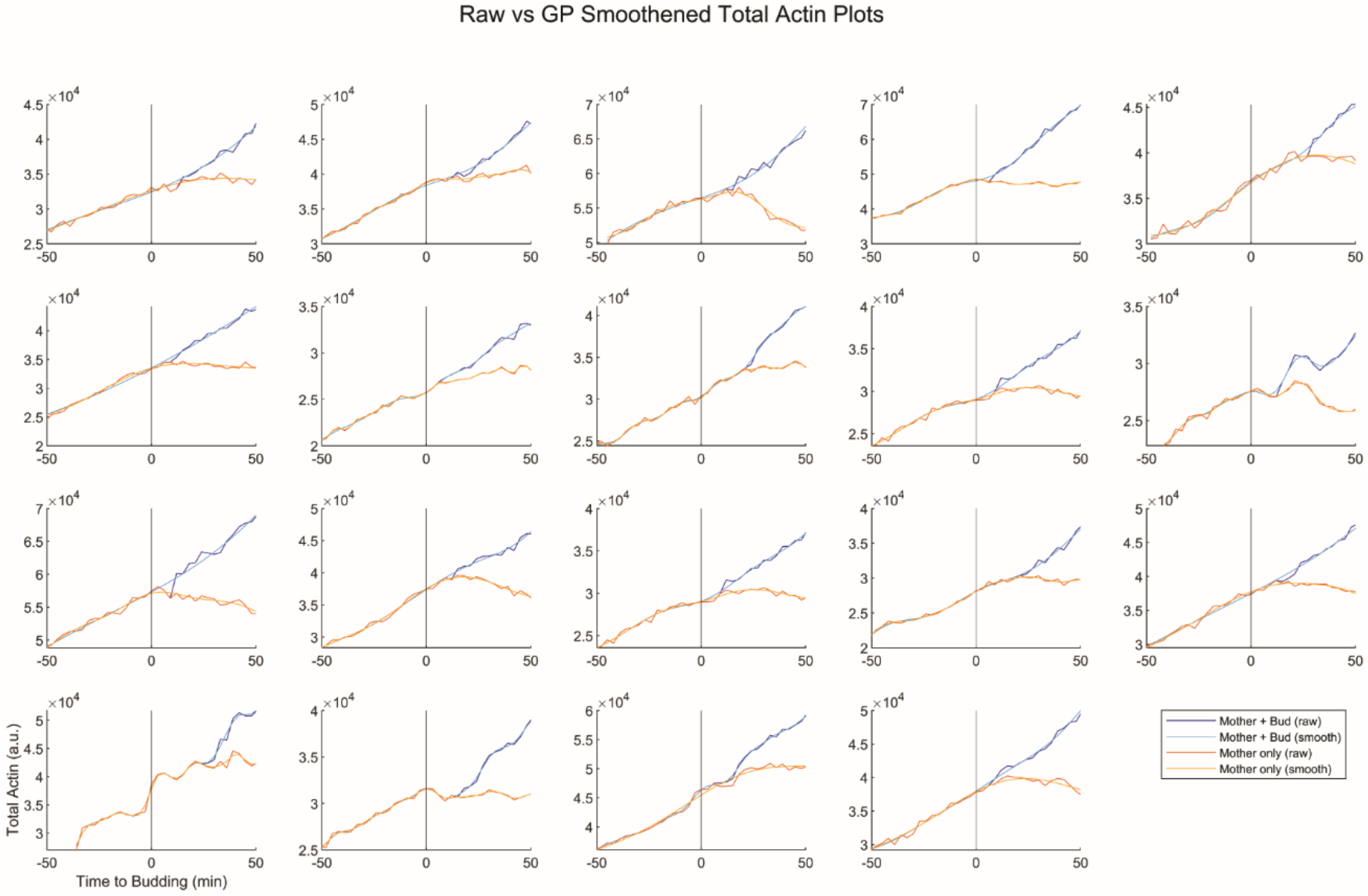
Single cell traces and fits for *ACT1pr-mCitrine* expressing strains. Examples of Gaussian process regression of data from Figure 3b). The total mCitrine data were smoothed using the Gaussian process regression performed in MATLAB using a pure quadratic basis function. The hyperparameters were optimized with at least 30 evaluations. Data from 19 random cells were chosen to be displayed to illustrate typical fits for single cells.

## Supplementary Information: Schneider lab

### Live-cell microscopy

#### Nikon Eclipse setup with custom microfluidics setup

Live-cell time-lapse microscopy was performed in a custom-made microfluidic device made of polydimethylsiloxane (PDMS) and a glass cover-slip that allows trapping of cells in a dedicated region of interest, limiting colony growth in the XY-plane. Cells were grown on synthetic complete media with 2% glycerol 1% ethanol as a carbon source (SCGE) overnight to exponential phase prior to the experiment. During the experiment, SCGE was supplied at a constant flow of 20 μl/min, enabling imaging of colony growth over several generations.

A Nikon Eclipse Ti-E with SPECTRA X light engine illumination and an Andor iXon Ultra 888 camera was used for epifluorescence microscopy. A plan-apo λ 60x/1.4Na Ph3 oil immersion objective was used to take phase contrast and fluorescence images with a 3-minute frame rate. For automated focusing, the built-in Nikon perfect focus system was used during the experiment. mCitrine fluorescence was imaged by exposure for 400 ms, illuminating with the SPECTRA X light engine at 504 nm and about 16 mW (25%) power.

Temperature control was achieved by setting both a custom made heatable insertion and an objective heater to 30°C.

Identical settings were used for each of the experiments.

### Image Analysis

#### Cell segmentation

Cells were automatically segmented based on phase contrast images using the *Matlab* based *Phylocell* software developed in the Gilles Charvin lab^20^. Segmentation results were visually inspected and manually corrected if necessary.

#### Calculation of cell volume

To calculate cell volume based on 2D phase contrast images, we first aligned the segmented cell area along its major axis. Next, we divided the cell into slices perpendicular to the major axis, each 1 pixel in width. To approximate cell volume, we then assumed rotational symmetry of each slice around its middle axis parallel to the cell’s major axis. We then calculated for each slice the volume of the resulting cylinder with 1 pixel in height and diameter the width of the respective slice. Finally, we summed the volumes of each cylinder to obtain total cell volume.

#### Protein amount calculation

To calculate changes in Whi5 amount over time, we used a strain where Whi5 was endogenously tagged with mCitrine. In this case, the total fluorescence signal produced by mCitrine should directly correlate with Whi5 amount. To measure the mCitrine signal, we first estimated the background fluorescence using two areas of 200×200 pixels containing no cells and subtracted its median value from all pixels. Then, we calculated the total fluorescence for each cell separately as a sum of intensities of all pixels within a cell counter.

#### Cell cycle analysis

We analyzed 100 cells for the Whi5-mCitrine strain and 59 cells for the wildtype control (KSY108-1 & MMY116-2C, respectively; Schmoller et al. 2015^2^) during their first cell cycle. Next, we estimated the median G1 beginning (birth time) as 69 minutes and the median time of cytokinesis as 96 minutes from the bud emergence. For cell cycle analyses shown in Fig. S2b, we divided each total fluorescence value of each cell by the corresponding cell volume to account for differences in the cell volume over time. For Supplementary Fig. S2b, we corrected for the wildtype autofluorescence by subtracting its mean value for each time point from values of Whi5-mCitrine strain for the same time points. Then, we additionally normalized each fluorescence value of each cell by the fluorescence value at birth. Finally, we pooled all data for Whi5-mCitrine, and plotted the mean fluorescence from the median G1 beginning (birth time) to bud emergence. For Fig. S2a, we pooled all data for the Whi5-mCitrine strain without any correction or normalization and plotted the mean fluorescence from the median G1 beginning (birth time) to the median time of cytokinesis. For both subpanels of Supplementary Fig. S2, 95% confidence intervals were determined from 50000 bootstrap samples.

### Re-analysis of mass spectrometry data

Our re-analysis of the mass spectrometry experiments performed by Litsios et al. utilized the .raw MS acquisition files and SpectroDive output files that were uploaded to the PRIDE repository (PXD015327). We calculated the approximate TIC for each experimental time point from the corresponding .raw data file (see “TIC” tab of Table S1). To quantify changes in the amount of yeast peptide from time point-to-time point, we used a corresponding DDA spectral library and the Skyline analysis software to extract the chromatograms for several hundred yeast peptides in each experimental time point (DDA acquisition files were provided upon request from the authors of Litsios et al.). Similar to what was performed by Listos et al., we selected a set of 6 peptides from 3 different yeast housekeeping proteins (Enolase, ADH1, and GAPDH) and used the average fluctuation in their peptide peak areas to approximate changes in the total amount of yeast protein within each sample (see “normalization” tab of Table S1). These peptides behaved similarly to many other yeast housekeeping genes and their peaks consistently traced to a similar retention time (see “normalization” tab of Table S1).

The supplemental file related to the PRM analysis that was directly uploaded with the Litsios et al. article (Supplemental Table 4 from Litsios et al.) contains the top seven Q3 transitions for each Cln3 and Whi5 peptide but no quantitative information. A set of text files uploaded to PRIDE (“PRIDE_experiment1_spectrodive” and “PRIDE_experiment2_spectrodive”) contained the Cln3 and Whi5 PRM quantitation acquired from a set of 4-6 Q3 transitions. To re-create the analysis from Litsios et al., we normalized their published PRM measurements of the Cln3 and Whi5 peptides to the MS1 peptide peak areas of 3 yeast housekeeping proteins we extracted using Skyline. Litsios et al. used a single Q3 transition (the most abundant one) to quantify Whi5 and Cln3 abundance. A step-by-step demonstration of our calculations is laid out in Table S1. The authors of Listos et al. confirmed the correspondence of our re-analysis with the data they used to generate the plots in their manuscript.

### Methods Aldea Lab

Time-lapse microscopy and measurements of volume growth rate in G1 were as described by Ferrezuelo et al.^21^. Photobleaching during acquisition was negligible (less than 0.1% per time point) and background autofluorescence was always subtracted. The cellular concentration of fluorescent fusion proteins was obtained by dividing the integrated fluorescence signal within the projected area of the cell by its volume.

### Methods Kellogg Lab

Cell size was measured using a Coulter counter (Channelizer Z2; Beckman Coulter) as previously described in Lucena *et al*. ^7^. The percentage of budded cells was measured by counting the number of small unbudded cells over a total of more than 200 cells using a Zeiss Axioskop 2 (Carl Zeiss) and an AxioCam HRm camera with a 63 x 1.4 numerical aperture objective. Densitometric quantification of Whi5 and Cln3 western blot signals was performed using ImageJ^22^ normalizing over the loading control band.

## DATA TRANSFER AGREEMENT

**Parties:**

1. The University of Groningen, for the purpose of the Faculty of Science and Engineering, Nijenborgh 9, 9747 AG, Groningen, The Netherlands, duly represented by Matthias Heinemann, Professor, hereafter the “UG”;
and
2. <Name of institution/university>, <address>, <city>, <country>, duly represented by <first name and last name>, <job title>, hereafter the “Recipient”; Hereafter jointly referred to as the “Parties” and individually as “Party”‘;

**whereas:**

1. Parties have agreed upon the transfer of a data set, managed by the UG, (hereafter the “Data”) to Recipient;
2. Parties wish to lay down their rights and obligations regarding the Data transfer in a written agreement;

**have agreed as follows:**

<u>Article 1. Object of the agreement</u>

1. The Data covered by this agreement include a. a data set containing time-lapse microscopy data for Whi5-sfGFP concentration in cells grown on glucose and the associated autofluorescence control, managed by Matthias Heinemann, b. any related know-how transferred by the UG to Recipient and any data that is derived from the data set or which could not have been produced but for the use of the data set.

<u>Article 2. Data property rights</u>

1. The Data and all intellectual property rights concerning the Data shall remain the sole and exclusive property of the UG and its licensors. The UG shall be considered free, in its sole discretion, to distribute the Data to others for any purposes and to use it for its own purposes.

<u>Article 3. Usage rights</u>

1. Recipient is entitled to use the Data in accordance with this agreement. Recipient shall use the Data only for the purpose of conducting not-for-profit scientific research. Specifically, the Recipient will <add detailed description of research/intended use of the data> (hereafter: the “Research”).
2. The Data may only be used under supervision of <first name and last name of the principal investigator of Recipient>, <job title>. The Data may only be shared with employees of the Recipient that contribute to the Research.
3. Recipient agrees that nothing in this agreement shall be deemed to grant any rights under any UG patent applications or patents or any rights to use the Data for any purpose other than not-for-profit research.
4. Recipient has no rights in the Data other than as provided in this agreement, and at the request of the UG, Recipient will return or destroy all copies of the Data.

<u>Article 4. Research results</u>

1. Recipient will inform the UG, in confidence, of Research results related to the Data.
2. If Recipient desires to publish about the Research, Recipient will notify the UG not less than sixty (60) days prior to submission of its draft publication to a publisher or to any third party. The UG has the right to review the publication and to suggest alterations and additions.
3. If publication results from research using the Data, Recipient agrees to acknowledge the UG and/or give credit to UG scientists, as scientifically appropriate, based on the use of the Data and, if applicable, any other contribution they may make to the Research.

<u>Article 5. Confidentiality</u>

1. Recipient shall keep the Data strictly confidential. Recipient shall not disclose the Data to any third party or otherwise use it for its own benefit or for the benefit of a third party, without first obtaining written consent from the UG.
2. If Recipient breaches its confidentiality obligations of this agreement, it shall, without any notice of default or judicial intervention being required, for the benefit of the UG forfeit an immediately payable penalty not susceptible to setoff of EUR 10.000,-- (ten thousand euros), increased by an amount of EUR 1.000,-- (thousand euros) for each day that the breach continues. This penalty shall be without prejudice to the UG’s right to claim, in addition to this penalty, performance or full damages.

<u>Article 6. Protection of personal data (optional clause)</u>

1. This article 6 is only applicable if the Data contain personal data as defined in article 4 (1) of the EU General Data Protection Regulation (hereafter: the “GDPR”).
2. Parties will comply with the GDPR and other applicable legislation and regulations concerning the processing of personal data. Parties will determine in good faith how they will apply these laws and the data processing principles cooperating within the Parties.
3. If Parties should be considered joint controllers as defined in article 26.1 of the GDPR, they will determine their respective responsibilities for compliance with the GDPR and their respective duties to provide data subjects with information about the processing of their personal data. Parties will do so by means of an arrangement in writing.
4. If the Data are transferred beyond the borders of the European Economic Area, Parties will implement the safeguards required by the GDPR regarding such transfer.

<u>Article 7. Information security</u>

1. UG shall disclose the Data to Recipient via a secure sharing mechanism to be selected by the UG, using an encrypted connection for transporting the Data.
2. Recipient will store the data in a secure environment. Recipient warrants that the Data will be protected with appropriate technical and organizational security measures. Recipient shall at least implement the following security measures:

a. |<add description of security measures_>|

<u>Article 8. Limitation of liability</u>

1. The Data are experimental in nature and the Data are provided without warranty of merchantability or fitness for a particular purpose or any other warranty, express or implied. The UG makes no representation or warranty that the use of the Data will not infringe any patent or other proprietary right.
2. In no event shall the UG be liable for any use by Recipient of the Data or any loss, claim, damage or liability, of whatsoever kind or nature, which may arise from or in connection with this agreement or the use of the Data. Recipient agrees to indemnify the UG, its trustees, officers, agents, and employees from any liability, loss or damage they may suffer as a result of claims, demands, costs or judgments against them arising out of the use or disposition of the Data by Recipient.

<u>Article 9. Miscellaneous</u>

1. Parties designate employees with whom the execution of the agreement is invested. UG employee Matthias Heinemann, Professor, is contact person for the Recipient. Recipient’s employee <name>, <job title> is contact person for the UG.
2. This agreement will enter into force as of the date both Parties have signed it. This agreement will continue to be effective during the period that the Recipient possesses the Data. During that period, Recipient cannot terminate this agreement.
3. The UG may terminate this agreement if Recipient has not cured a breach of this agreement within a reasonable period of being notified of such breach.
4. None of the terms of this agreement shall be amended or modified except in writing and signed by the Parties hereto.
5. Parties are and intend to remain independent contractors. Nothing in this Agreement shall be construed as an agency, joint venture or partnership between the Parties.
6. This Agreement is not assignable by Recipient, whether by operation of law or otherwise, without the prior written consent of the UG.
7. This Agreement shall be governed by Dutch law only, without recourse to its conflict of law principles. Any dispute arising under this Agreement shall be settled by the district court in Groningen, the Netherlands.

**In witness whereof:**

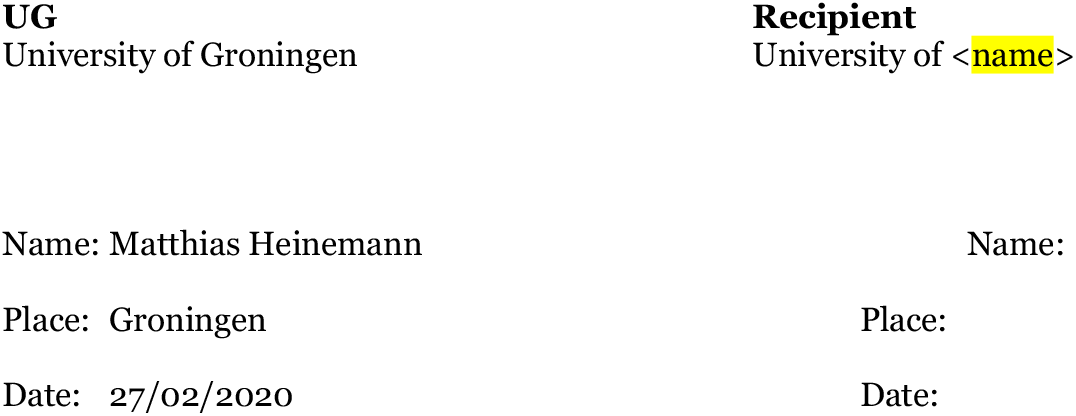

## Notes

### Competing Interest Statement

The authors have declared no competing interest.

## References

1. Litsios, A. et al. Differential scaling between G1 protein production and cell size dynamics promotes commitment to the cell division cycle in budding yeast. Nat. Cell Biol. 21, 1382–1392 (2019).

2. Schmoller, K. M., Turner, J. J., Kõivomägi, M. & Skotheim, J. M. Dilution of the cell cycle inhibitor Whi5 controls budding yeast cell size. Nature 526, 268–272 (2015).

3. Spellman, P. T. et al. Comprehensive Identification of Cell Cycle–regulated Genes of the Yeast *Saccharomyces cerevisiae* by Microarray Hybridization. Mol. Biol. Cell 9, 3273–3297 (1998).

4. Pramila, T., Wu, W., Miles, S., Noble, W. S. & Breeden, L. L. The Forkhead transcription factor Hcm1 regulates chromosome segregation genes and fills the S-phase gap in the transcriptional circuitry of the cell cycle. Genes Dev. 20, 2266–78 (2006).

5. Granovskaia, M. V. et al. High-resolution transcription atlas of the mitotic cell cycle in budding yeast. Genome Biol. 11, (2010).

6. Gomar-Alba, M., Méndez, E., Quilis, I., Bañó, M. C. & Igual, J. C. Whi7 is an unstable cell-cycle repressor of the Start transcriptional program. Nat. Commun. 8, 329 (2017).

7. Lucena, R. et al. Cell Size and Growth Rate Are Modulated by TORC2-Dependent Signals. Curr. Biol. 28, 196–210.e4 (2018).

8. Qu, Y. et al. Cell Cycle Inhibitor Whi5 Records Environmental Information to Coordinate Growth and Division in Yeast. Cell Rep. 29, 987–994.e5 (2019).

9. Barber, F., Amir, A. & Murray, A. W. Cell size regulation in budding yeast does not depend on linear accumulation of Whi5. bioRxiv (2020).

10. Chen, Y., Zhao, G., Zahumensky, J., Honey, S. & Futcher, B. Differential Scaling of Gene Expression with Cell Size May Explain Size Control in Budding Yeast. Mol. Cell 1–12 (2020). doi:10.1016/j.molcel.2020.03.012

11. Dorsey, S. et al. G1/S Transcription Factor Copy Number Is a Growth-Dependent Determinant of Cell Cycle Commitment in Yeast. Cell Syst. 6, 539–554.e11 (2018).

12. De Bruin, R. A. M., McDonald, W. H., Kalashnikova, T. I., Yates, J. & Wittenberg, C. Cln3 activates G1-specific transcription via phosphorylation of the SBF bound repressor Whi5. Cell 117, 887–898 (2004).

13. Costanzo, M. et al. CDK activity antagonizes Whi5, an inhibitor of G1/S transcription in yeast. Cell 117, 899–913 (2004).

14. Wagner, M. V. et al. Whi5 regulation by site specific CDK-phosphorylation in Saccharomyces cerevisiae. PLoS One 4, (2009).

15. Doncic, A., Falleur-Fettig, M. & Skotheim, J. M. Distinct Interactions Select and Maintain a Specific Cell Fate. Mol. Cell 43, 528–539 (2011).

16. Jorgensen, P. et al. The size of the nucleus increases as yeast cells grow. Mol. Biol. Cell 18, 3523–3532 (2007).

17. Walters, A. D. et al. The yeast polo kinase Cdc5 regulates the shape of the mitotic nucleus. Curr. Biol. 24, 2861–2867 (2014).

18. Chandler-Brown, D., Schmoller, K. M., Winetraub, Y. & Skotheim, J. M. The Adder Phenomenon Emerges from Independent Control of Pre- and Post-Start Phases of the Budding Yeast Cell Cycle. Curr. Biol. 27, 2774–2783.e3 (2017).

19. Elliott, S. G. & McLaughlin, C. S. Rate of macromolecular synthesis through the cell cycle of the yeast Saccharomyces cerevisiae. Proc. Natl. Acad. Sci. 75, 4384–4388 (1978).

20. Goulev, Y. et al. Nonlinear feedback drives homeostatic plasticity in H2O2 stress response. Elife 6, e23971 (2017).

21. Ferrezuelo, F. et al. The critical size is set at a single-cell level by growth rate to attain homeostasis and adaptation. Nat. Commun. 3, 1012 (2012).

22. Schneider, C. A., Rasband, W. S. & Eliceiri, K. W. NIH Image to ImageJ: 25 years of image analysis. Nat. Methods 9, 671–675 (2012).

